# Lox’d in translation: Contradictions in the nomenclature surrounding common lox site mutants and their implications in experiments

**DOI:** 10.1101/2020.07.14.202044

**Authors:** Daniel Shaw, Luis Serrano, Maria Lluch-Senar

## Abstract

The Cre-Lox system is a highly versatile and powerful DNA recombinase mechanism, mainly used in genetic engineering to insert or remove desired DNA sequences. It is widely utilised across multiple fields of biology, with applications ranging from plants, to mammals, to microbes. A key feature of this system is its ability to allow recombination between mutant lox sites, traditionally named lox66 and lox71, to create a functionally inactive double mutant lox72 site. However, a large portion of the published literature has incorrectly annotated these mutant lox sites, which in turn can lead to difficulties in replication of methods, design of proper vectors, and confusion over the proper nomenclature. Here, we demonstrate common errors in annotations, the impacts they can have on experimental viability, and a standardised naming convention. We also show an example of how this incorrect annotation can induce toxic effects in bacteria that lack optimal DNA repair systems, exemplified by *Mycoplasma pneumoniae*.

**Data Summary:** The authors confirm all supporting data, code and protocols have been provided within the article or through supplementary data files.

## Introduction

The Cre-Lox system was first characterised in 1981 by Nat Sternberg and Daniel Harrison (1). It is a DNA recombinase system derived from the P1 bacteriophage, and consists of two components. The first, the Cre recombinase, is a 38-kDa monomeric tyrosine recombinase. This protein acts upon a pair of 34bp lox sites (**<u>l</u>**ocus **<u>o</u>**f (**<u>x</u>**)crossing over), which consist of an 8-bp central spacer region flanked by two 13-bp inverted repeat regions (2,3). The central spacer unit of the lox site gives it a ‘directionality’, which dictates the outcome of the reaction between the Cre recombinase and DNA, as shown in Figure 1. This outcome depends on the orientation of the mutant lox sites. If they are in the same (*cis*) orientation, recombination results in the circularisation of the genomic DNA between the two lox sites, and thus its removal from the genome. If the lox sites are in opposite (*trans*) orientations, then the DNA between them is inverted as a consequence of the recombination event.

**Figure 1:**
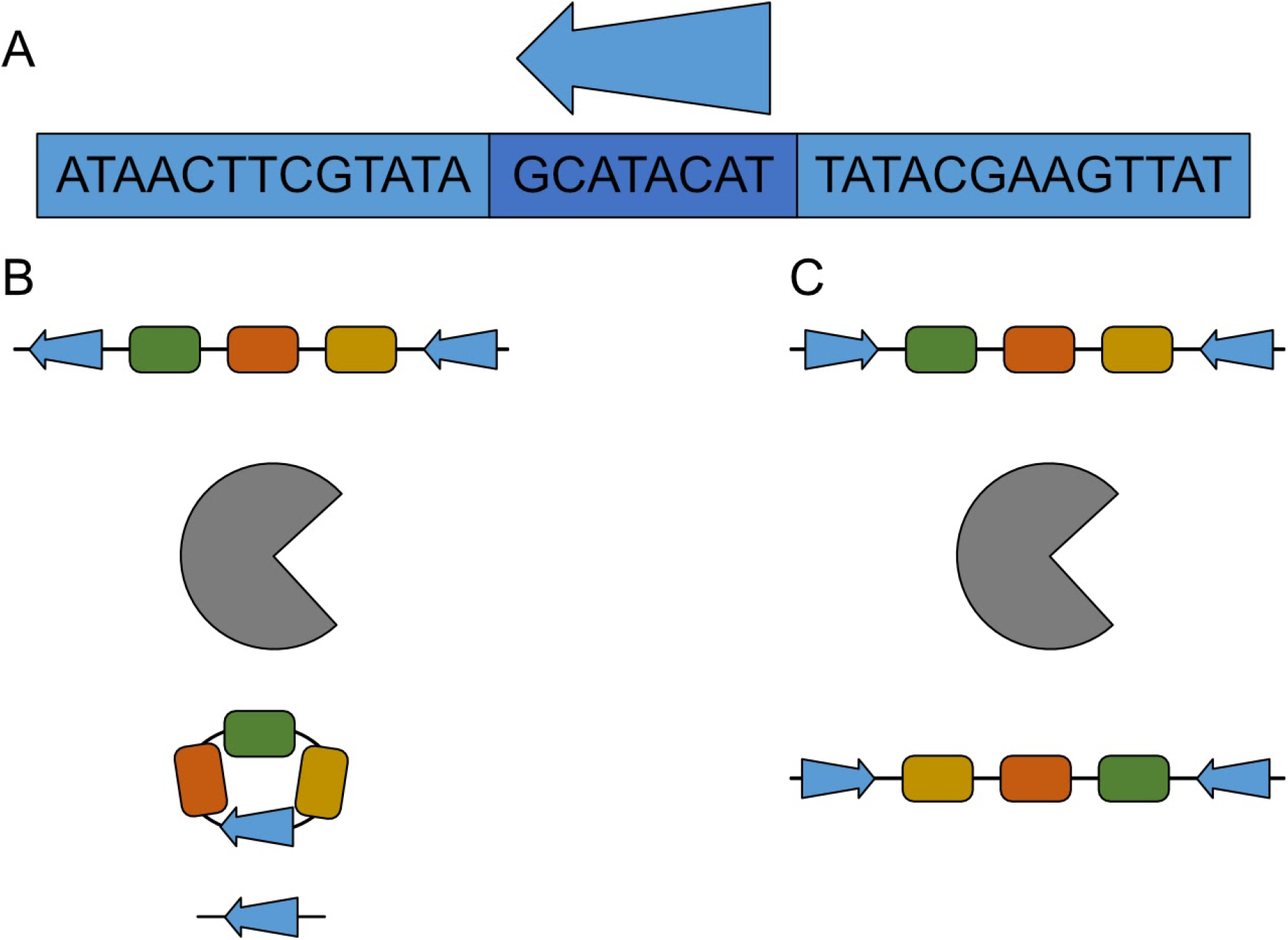
Sequence and orientation of the WT loxP site, and effect of orientation on lox recombination reactions. (A) Sequence of the loxP site, with the 13-bp inverted repeats in lighter blue, and the central spacer region in darker blue. The arrow indicates the directionality of the lox site. (B) Action of the Cre recombinase (grey) on two lox sites in *cis* orientation. DNA between the two sites is excised and circularised, leaving 1 lox site in the genome and 1 lox site on the circular DNA. (C) Action of the Cre recombinase when lox sites are in *trans*. DNA between the two sites is inverted.

For the sake of brevity, a full review into the history of the Cre recombinase and its reaction kinetics are outside the scope of this paper, and have been discussed at detail by other authors (4–7). Instead, here we will focus on one aspect of the Cre-Lox system; its ability to utilise mutant lox sites, and how these sites interact.

The two most widely used lox mutations are the lox66 and lox71 sites, which were first described in 1995 (8). These lox mutants contain alterations to their initial or terminal 5 base pairs, with the lox66 having the first 5 bases changed, and lox71 having the last 5 bases changed (in regard to the canonical loxP), shown below in Table 1:

**Table 1:**
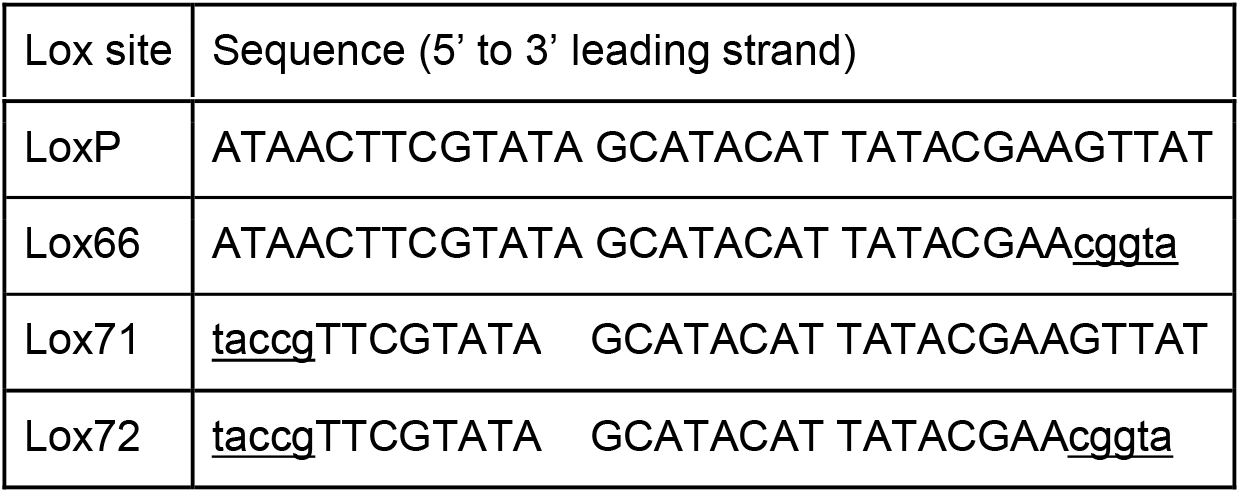
The sequence of the canonical loxP site and its mutants, with directionality of the lox site from left to right.

The Cre recombinase exists as a monomer, however during recombination it forms a tetramer, with one molecule attached to each flanking region of the two lox sites (5). As shown in Figure 2 below, modified from the review of the Cre recombinase by Gregory Van Duyne (5), the Cre complex manoeuvres the two DNA strands so that the lox sites are brought together. The arrows indicate the directionality of the lox sites, and as the complex is formed they face in opposite directions. The lagging strand from both DNA molecules is then cleaved, and annealed to the other DNA molecule via a Holliday junction-forming reaction. The same then occurs for the other strand, resulting in the left element of the first lox site becoming joined to the right element of the second, and vice versa.

**Figure 2:**
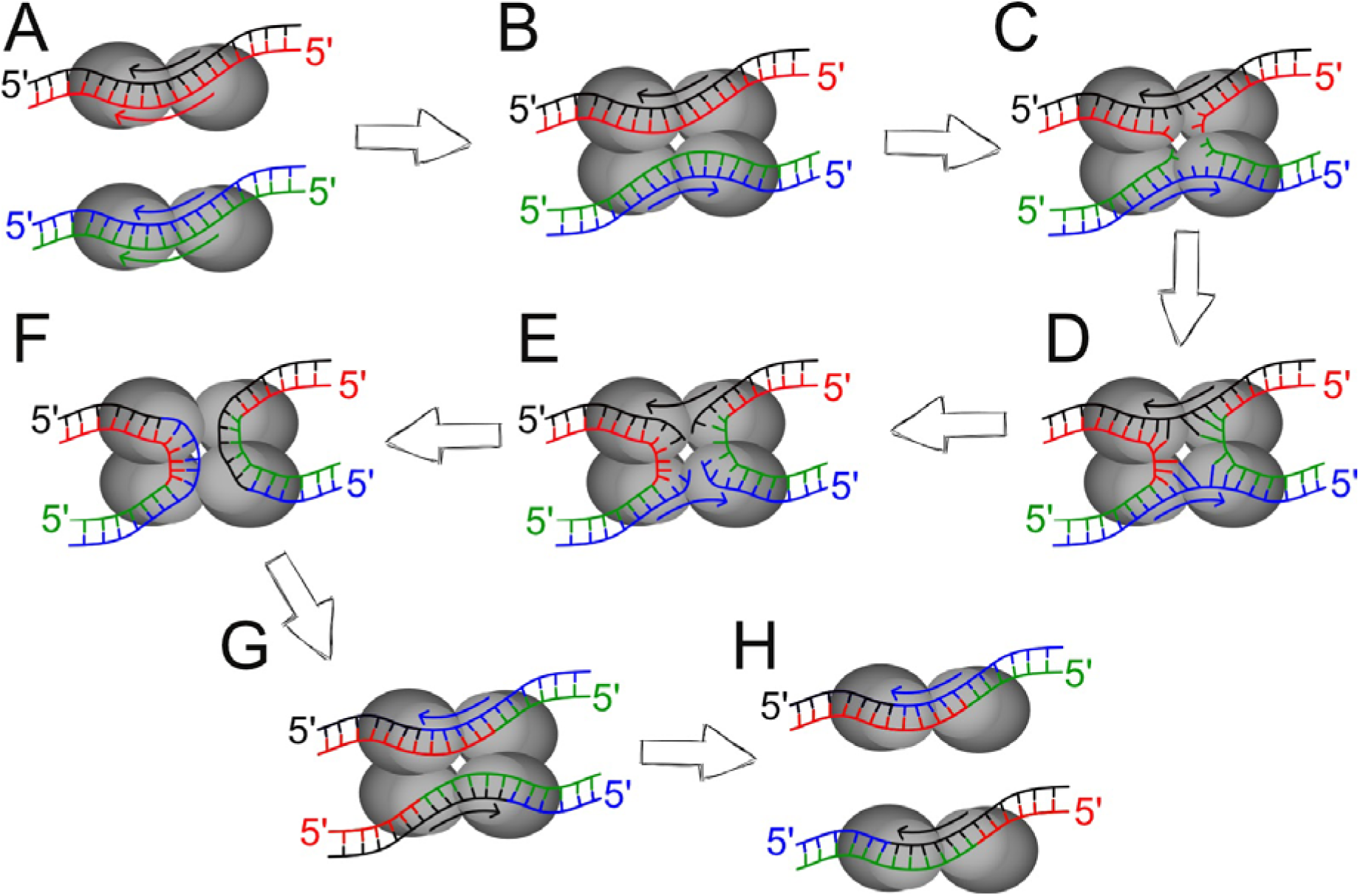
Modified from Van Duyne (5). Shows the reaction pathway of the Cre recombinase complex on two loxP sites, with the stands of each DNA molecule shown in separate colours, with the top strands in black and blue, and the bottom strands in red and green. The direction of the lox sites is indicated by the coloured arrows. (A) The Cre recombinase molecules bind to each homology arm of each lox site (B) The Cre-lox complex is formed, with the four Cre molecules bringing the DNA together so the two lox sites run antiparallel to each other. (C) Cleavage of the bottom strands by Tyr324 in the Cre subunits. (D) Formation of 3’-phosphotyrosine linkages between the bottom strands leads to the creation of a 4-way Holliday junction between the two DNA molecules. (E) Cleavage of the top strands allows for strand exchange between the two DNA molecules. (F) Top strands of both molecules are ligated together. (G) Formation of the two new lox sites. (H) New lox sites contain the 5’ homology arm of the first lox site, with the 3’ homology arm of the second, and *vice versa*.

As both lox sites in the diagram are identical, they do not change their sequence via recombination, as every flanking region is identical. However, as each new lox site generated is a combination of both previous sites, when two different mutant lox sites are incorporated, a double mutant is always created. This is due to the Cre recombinase complex to manoeuvre the DNA strands into an antiparallel formation ensures that one ‘side’ of the reaction will always contain both mutant flanking regions, and thus will combine together (9).

The Cre recombinase can recognise the single mutant lox66 and lox71 with the same affinity as the loxP, however the double mutant lox72 shows a marked decrease in affinity for the Cre (8), making the site functionally silent. This lack of affinity for the lox72 site allows for multiple lox reactions to occur in a genome in series, without affecting or cross-reacting with each other (10–12).

In this paper, we show a comprehensive analysis of the literature associated with lox66 and lox71 sites, with examples of the common errors that have occurred in the nomenclature and a standardised naming convention to help resolve these remaining ambiguities. We also highlight the dangers of such errors, demonstrating that the presence of a single active lox site within a bacterial genome can initiate a lethal phenotype, on par with a double strand break in the DNA. We use *Mycoplasma pneumoniae* as a model organism, and demonstrate that the lethality associated with the Cre acting on a single lox is analogous to a double stranded break generated by the meganuclease *I-SceI*, which we also report to be active in *M. pneumoniae* for the first time.

With the demonstration that there occurs mis-annotation of both the mutant arms of the lox sites, and also their directionality, we demonstrate the potentially destructive consequences of when these errors are compounded. This occurred within our own lab, and we hope that this publication may avert the same issues in future experiments, and potentially explain why some previous experiments may have failed.

## Methods

### Literature review of lox site nomenclature

A literature review aiming to capture as many of the papers describing the interactions between lox66 and lox71 sites was performed (See Figure 3). Many papers were present via multiple search terms, so replicates were removed. The papers were then queried for their relevance to the topic, specifically if they included the use of the lox66 or lox71 site within their methodology or discussion. Those papers that used the sites in a material way were included, and all others discarded.

**Figure 3:**
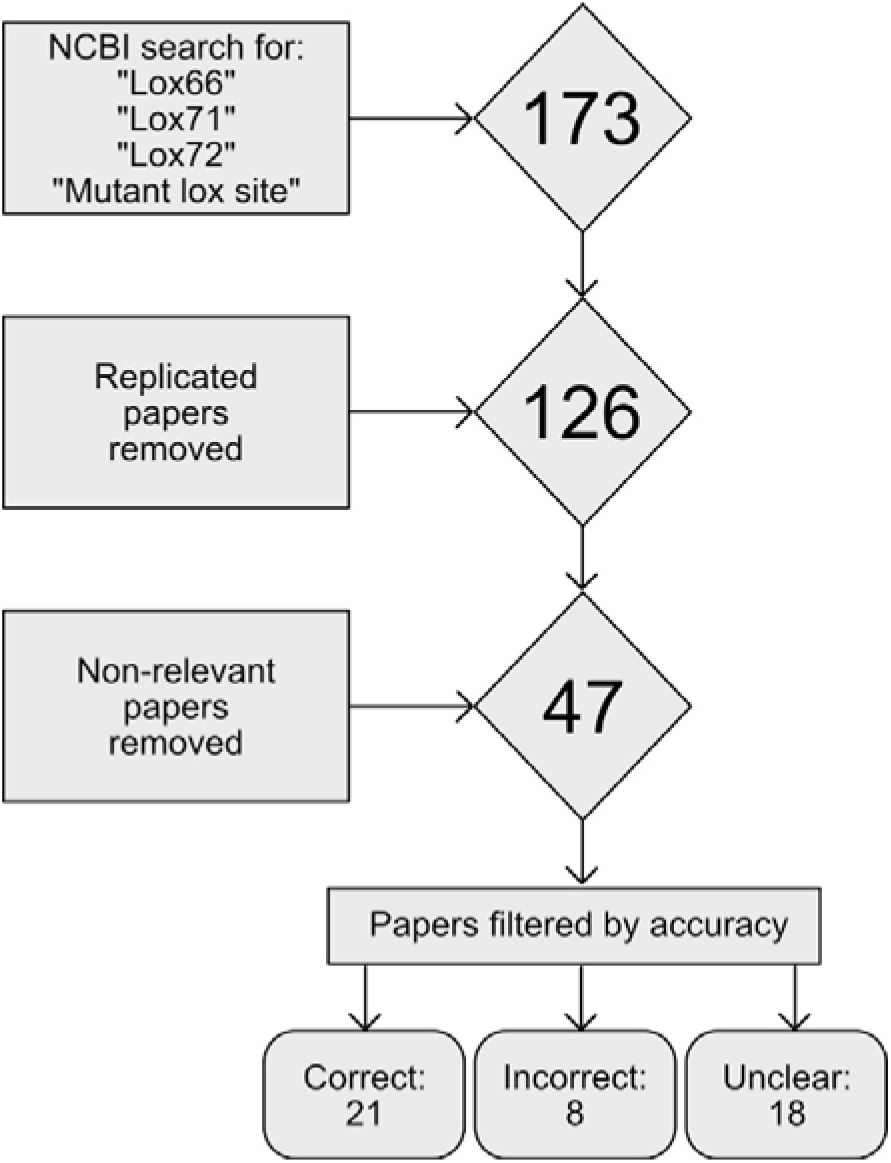
Flowchart describing the generation of the literature review. The square boxes show the data acquisition and filtering steps in order of occurrence, and the diamonds show the number of papers being considered after each step. The final 47 chosen papers were analysed for their use of lox nomenclature, and classified appropriately, as described in the text above.

The remaining 47 papers were then judged for their accuracy in regard to the lox site nomenclature. The benchmark used for accuracy was the nomenclature described in the original paper by Albert et al., (8). Papers which provided the sequence of the lox sites they used were filtered into either the “Correct” or “Incorrect” category, depending on if the nomenclature they used was in concordance with Albert et al. All papers that did not show the sequence of the lox sites were placed within the “Unclear” category.

### Strains and culture conditions

Wild-type *Mycoplasma pneumoniae* strain M129 (ATTC 29342, subtype 1, broth passage no. 35) was used. Cells were cultured in 75cm^3^ tissue culture flasks at 37°C in standard Hayflick media, as described by Hayflick (13) and Yus et al., (14), supplemented with 100 μg/ml ampicillin, 2 μg/ml tetracycline, 20 μg/ml chloramphenicol, 3.3μg/ml puromycin and 200 μg/ml gentamicin as appropriate. Hayflick agar plates were created by supplementing the Hayflick with 1% Bacto Agar (BD, Cat. No. 214010) pre autoclave.

NEB 5-alpha Competent *E. coli* cells (New England Biolabs, Catalogue number C2987H) were used for plasmid amplification and cloning. They were grown at 37°C in Lysogeny Broth (LB) at 200RPM or static on LB agar plates, supplemented with 100μg/ml ampicillin.

### Plasmid DNA

All plasmids were generated using the Gibson isothermal assembly method (15). DNA was isolated from NEB 5-alpha Competent *E. coli* cells, and individual clones were selected for using LB + ampicillin plates (100 μg/ml). Correct ligation was confirmed via Sanger sequencing (Eurofins Genomics). A list of all plasmids, and primers used in their generation and sequencing can be found in Suppl. Tables 1 & 2.

### Single lox site lethality

WT M129 cells were transformed according to the protocol outlined by Hedreyda et al (16). Cells were grown to mid-log phase, identified by the Hayflick media changing from red to orange. The media was decanted and the flask was washed 3x with 10ml chilled electroporation buffer (EB: 8mM HEPES, 272nM sucrose, pH 7.4). Cells were scraped into 500μl chilled EB and homogenised via 10x passages through a 25-gauge syringe needle. Aliquots of 50μl of the homogenised cells were mixed with a pre-chilled 30μl EB solution containing the 1pMole of pMTn_Lox66_Sce_Cm plasmid DNA. Samples were then kept on ice for 15 mins. Electroporation was done using a Bio-Rad Gene Pulser set to 1250 V, 25 μF and 100Ω. After electroporation, cells were incubated on ice for 15 mins, then recovered into a total of 500μl Hayflick media and incubated at 37°C for 4 hours. 125μl of transformed cells were then inoculated into T75 culture flasks containing 20ml Hayflick and supplemented with 20 μg/ml chloramphenicol.

The transformed cells were grown to mid-log phase, then isolated. To ensure the recovery of planktonic cells, the media was transferred to a 50ml Falcon tube. The flask was then scraped into 500μl Hayflick, which was added to the Falcon tube with the media. The sample was centrifuged at 10,000RPM at 4°C for 10 mins to pellet the cells. The supernatant was discarded and the cells re-suspended in 500μl chilled EB. A fresh batch of WT M129 cells was also prepared. The cells were homogenised via 10x passages through a 25-gauge syringe needle. Aliquots of 50μl of the homogenised cells from both conditions were incubated with a pre-chilled 30μl EB solution containing the 1pMole of either the pBSK_p438_Sce_Puro, pBSK_p438_Cre_Puro or pBSK_p438_Puro plasmid, and transformed using the previously described settings. Cells were recovered in the same manner, and 125μl were inoculated into a T75 flask containing 20ml Hayflick media supplemented with XX μg/ml puromycin, and incubated at 37°C for 5 days.

After the incubation period, cells were isolated via the centrifugation protocol specified above. The pellets were dissolved in 500μl of fresh Hayflick media, and serial diluted. 10μl spots of each dilution were then plated on Hayflick agar plates, and incubated at 37°C for 1 week before being counted.

## Results

A large number of papers were omitted from the initial pool of papers, are two main reasons for this. First, all of the papers found with the searches for specific lox sites were also found in the “Mutant lox site” search, indicating that the remaining papers were more tangentially related to the topic. While almost all papers from the first three searches were relevant, very few papers from the final search were relevant that had not already been accounted for. Second, the “Mutant lox site” search returned a large number of papers that were not germane to the topic of this review. They generally fit into three further categories; i) Papers involving mutant lox sites that were not lox66 or lox71. ii) Papers describing mutants created by protocols using other lox sites, typically loxP or loxM variants. iii) Papers involving LOX genes, typically from *Pseudomonas aeruginosa*.

**Table 2:** Classification of the 47 relevant papers

| Correct papers | Incorrect papers | Unclear papers |
| --- | --- | --- |
| References (8,10,12,17–34) | References (35–42) | References 36–53 |

In total, we found that 8/47 (17%) of papers surveyed had incorrectly labelled their lox sites. Of these, 7 of the 8 papers (35,36,38–42) had simply mislabelled the lox sites, ascribing the lox66 name to the canonical lox71, and *vice versa*. The remaining paper (37) correctly annotated the lox sites, but implied the directionality of the lox site was in the opposite direction to which it was written.

The strict criteria employed in classifying papers as either correct or incorrect meant that a large number of relevant papers did not fit into either the correct or incorrect category. The main reason for this was that they did not explicitly state the sequences of the lox sites that they used. Looking at the figures and methodologies, all of them seem to follow convention and appear to use the lox sites correctly. Furthermore, almost every paper references either Albert et al., (8), Leibig et al., (24) or Araki et al., (17) when discussing the addition of lox sites into a construct. All three of these papers show correct annotations for the lox sites that they use.

However, just having the correct reference is not enough to guarantee that the lox site is correct. In Table 1 of Missirlis et al., (40), they show incorrect sequences for lox66 & lox71 while referencing Albert et al. (8) and Araki et al. (17) for both. Therefore, it seemed reasonable to separate the clear-cut cases from the ambiguous ones, as to create a ‘gold standard’ of reporting in relation to lox site nomenclature.

### Lethality conferred by single lox sites

We were interested in examining the potential effects of mis-labelling the lox sites on experimental viability. If one or more lox sites are not correct, or are not internally consistent, then there is the possibility of an active lox site being formed instead of a lox72, likely due to an inversion between the lox sites taking place instead of a deletion. The most damaging scenario would be both the mis-annotation of the lox sites, and their directionality. This occurred within our own laboratory, and resulted in months of troubleshooting before its resolution.

This is not encountered in the literature, though this may be via experiments failing and not generating publishable results. As such, we decided to test the effect of a single lox site being present in the genome of a bacteria with the Cre recombinase being expressed, to validate if this annotation error had caused the failure of the experiment.

Using *M. pneumoniae* as a model system, we transformed the cells with a transposon containing a single lox66 site, along with an *I-SceI* meganuclease recognition sequence. The *I-SceI* meganuclease induces a double stranded break in the genome, which is lethal to *M. pneumoniae* due to its limited DNA repair mechanisms (60), and provides a good baseline control for lethality. While the usage of the *I-SceI* meganuclease has not been previously demonstrated in *M. pneumoniae*, it has been expressed in other mollicute species, such as *Spiroplasma citri* (61), where it was also used to induce a lethal phenotype.

The *M. pneumoniae* cells were transformed with a transposon containing the *I-SceI* meganuclease recognition site, a lox66 site, and a chloramphenicol resistance gene. They were then transformed with a suicide vector containing either a Cre recombinase gene, the *I-SceI* meganuclease, or an empty control, alongside the WT cells.

As shown in Figure 4, there is a clear lethal phenotype exhibited by the action of the Cre on cell lines with a single lox site present, as after 5 days of incubation with the relevant plasmids, we demonstrate a 3-fold reduction in cell viability measured by CFU, which is near identical to the lethality induced by the double stranded break enacted by the *I-SceI*. We therefore demonstrate that both the action of the Cre on a single lox site can cause cell death in *M. pneumoniae*, and also that the *I-SceI* meganuclease can be expressed and active within the system.

**Figure 4:**
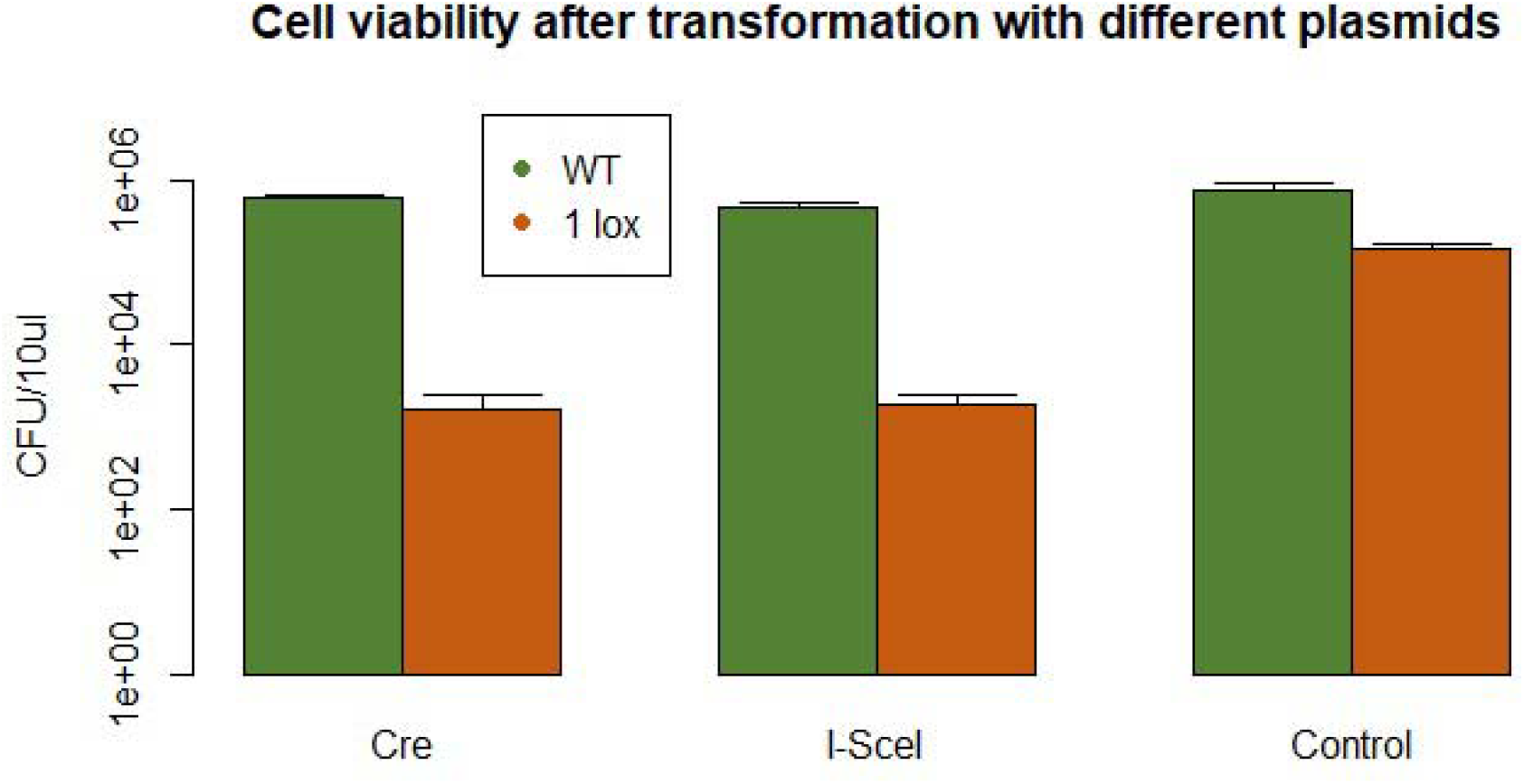
Effect of incorporating incorrect lox sites on experimental viability. CFU (colony forming unit) count after transforming a *M. pneumoniae* M129 containing a single lox site and a *M. pneumoniae* M129 WT strain with different suicide vectors containing either the Cre recombinase, *I-SceI* meganuclease or an empty control. Error bars indicate 2 standard deviations from the mean. Data from WT M129 cells shown in green, and M129 with a single lox site integrated into the genome in orange.

## Discussion

We have shown that there are a large number of errors found within the relevant lox literature. The main cause of these errors potentially stems from the reporting of the original sequences in the paper by Albert et al (8). In Figure 3 of their paper, the sequences are shown of the relevant lox sites. Here, the directionality of the lox is facing right to left. However, many of the papers with an error depict the lox site running left to right. In essence, they have switched the directionality of the central spacer region, without also changing which flanking arm contains a deletion.

**Figure 5:**
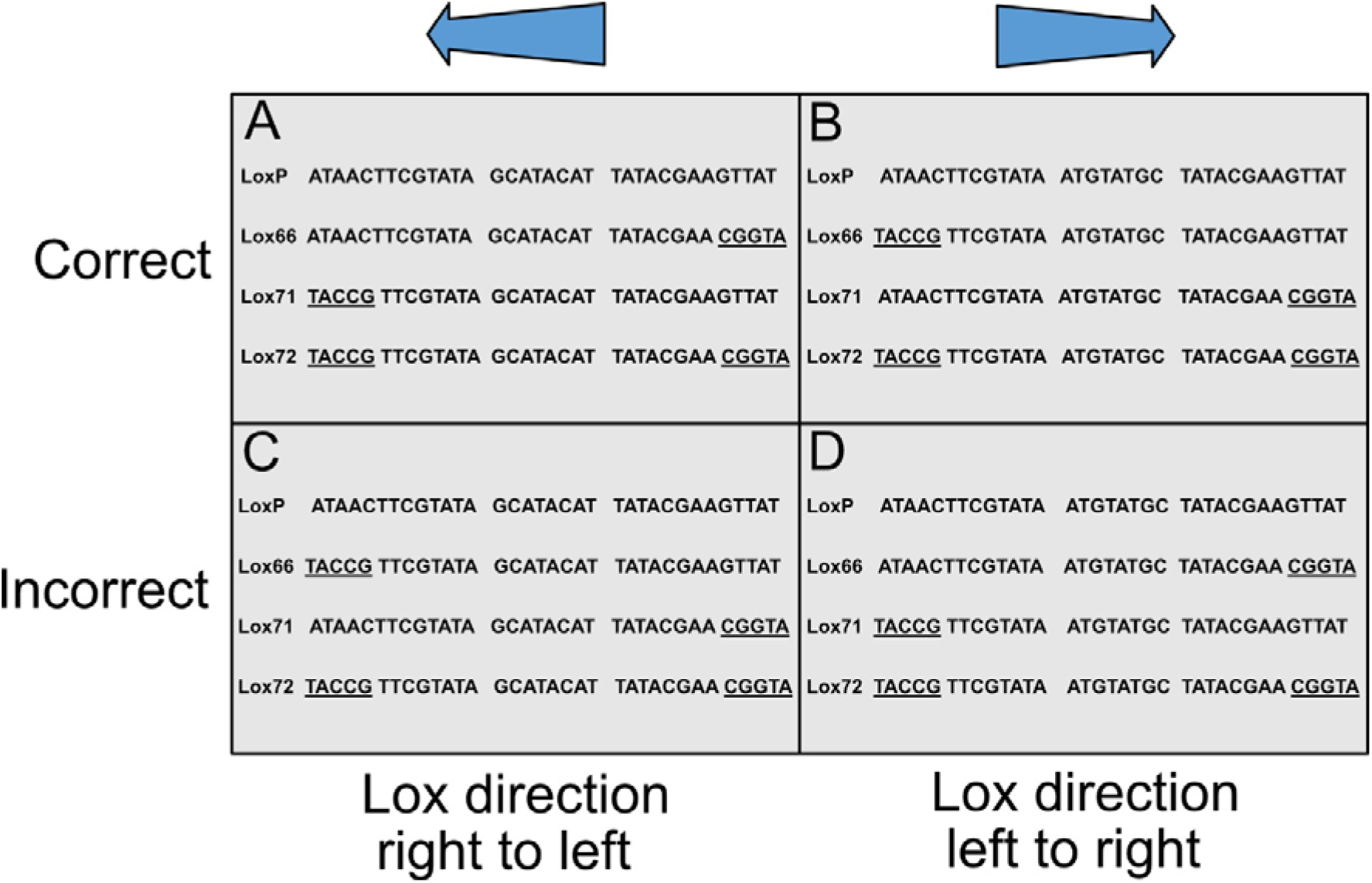
Common examples of lox mis-annotation, showing correct and incorrect examples for each directionality. All mutant regions are underlined. (A) Taken from Albert et al. (8). Shows the original description of the lox66, lox71 & lox72 in relation to the loxP. (B) Sequences taken from Leibig et al. (24). (C) Sequences taken from Weng et al. (42). (D) Sequences taken from Guan et al. (39).

In the original paper, Albert et al. (8) describes the sites as both lox66 & lox71, but also as “a site with a mutation in the [left element] (LE mutant site) and a site with a mutation in the [right element] (RE mutant site)”. However, the paper does not openly state which is which. In turn the reader is forced to interpret this as either left or right with regard to the directionality of the lox site, or 5’ to 3’ of the given sequence. Therefore, while the diagram of the mutant lox sites is clearly demarcated, the addition of the poorly characterized LE and RE annotations is ambiguous, and may have led to the errors described below.

It is worth emphasising at this point that these errors do not affect the scientific validity of the papers they are found in, as the constructs each author describes will create inactive lox sites. It is simply that the annotations the authors attribute the lox sites they use are incorrect. In the papers found with errors in lox site annotations, they fall into two distinct categories; i) miss-annotation of the lox site name, and ii) miss-annotation of the direction of the lox site.

A good example of the lox sites being simply miss-annotated is shown by Carter & Delneri (35). In their paper, they refer to the lox sites as Left Element and Right Element mutants to try and avoid confusion. Their premise is to flank genes for deletion with mutant lox sites, induce a deletion and create a silent lox72 site, as shown in Figure 1 of their paper, reproduced below:

As shown in Figure 6, the lox sites are inserted running left to right, as indicated by the arrows, with the 5’ or left element of the first lox mutated and the 3’ right element of the second lox mutated. This will give the mutant lox site shown in the figure. However, when looking at the table of oligos provided in Table 1 of the paper, reproduced below, the oligos used to create the lox sites do not match with their role in Figure 6.

**Figure 6:**
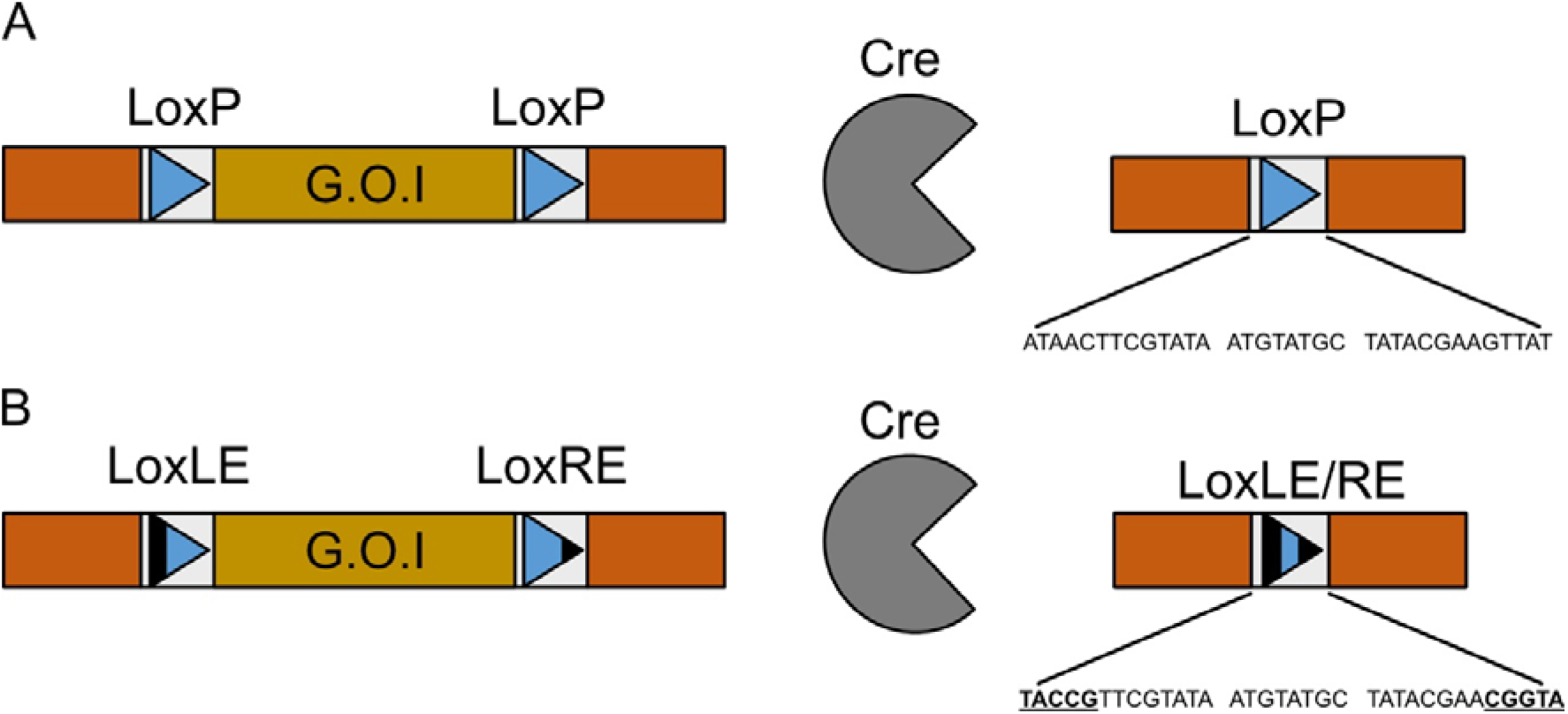
Modified version of Figure 1 from Carter and Delneri (35) showing the schematic of their Cre-lox deletion protocol, where their gene of interest (G.O.I) is flanked by different lox sites. (A) The action of the Cre on the DNA causes a deletion between the two LoxP sites, leaving a LoxP in the genome. (B) The action of the Cre on the DNA leaves an inactive double mutant lox site.

**Table 3:**
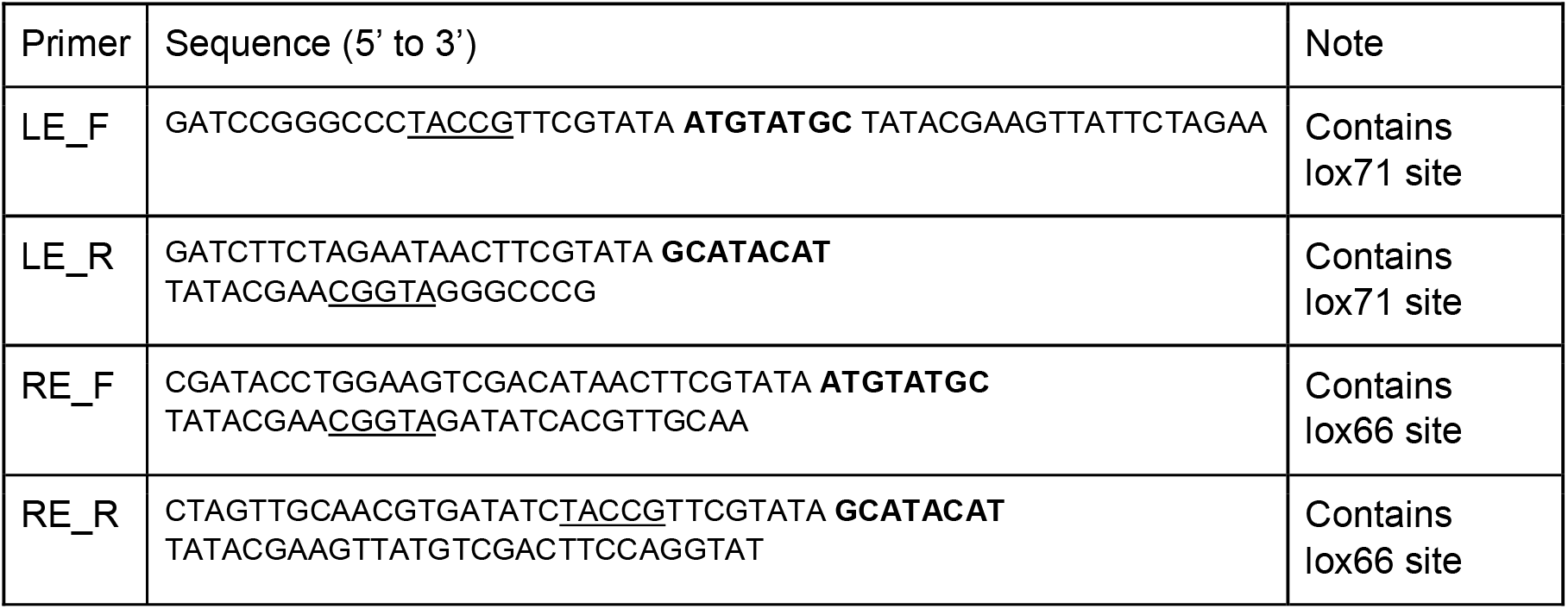
Modification of Table 1 from Carter and Delneri (35). Annotations to the sequences added for clarity, with spacer unit in **bold**,and the mutant regions <u>underlined</u>.

The “Note” column shows the authors’ annotations for the lox sites. However, both are incorrect, as when the lox site is written in a left to right directionality, the left arm mutant is the lox66, not the lox71 (24). The lox sites are still in the correct orientation, and the correct arms are mutated in each instance. Therefore, while the reaction schematic provided by Carter & Delneri is correct, and will give rise to a mutant lox72 site, the original lox sites are simply miss-labelled.

An example of attributing the wrong direction to the lox site can be found with Weng et al. (42). Figure 7 is an adaptation of Figure 1 from their paper. It shows a self-existing construct flanked by mutant lox sites, with their sequences shown below. The figure shows the lox sites in a left to right orientation, however the sequences of the lox sites shown below it are in a right to left direction. Therefore, the lox66 and lox71 annotations are incorrect. However, this will not affect the validity of the experimental design, as a lox72 will still be formed upon the action of the Cre. The lox66 site described as being in a left to right orientation is in truth a lox71 site in right to left orientation, with the opposite being true for the lox71 site. Simply, the arrows in the figure should be shown in the opposite direction, with the mutant regions of the lox sites still on the 5’ and 3’ arms respectively.

**Figure 7:**
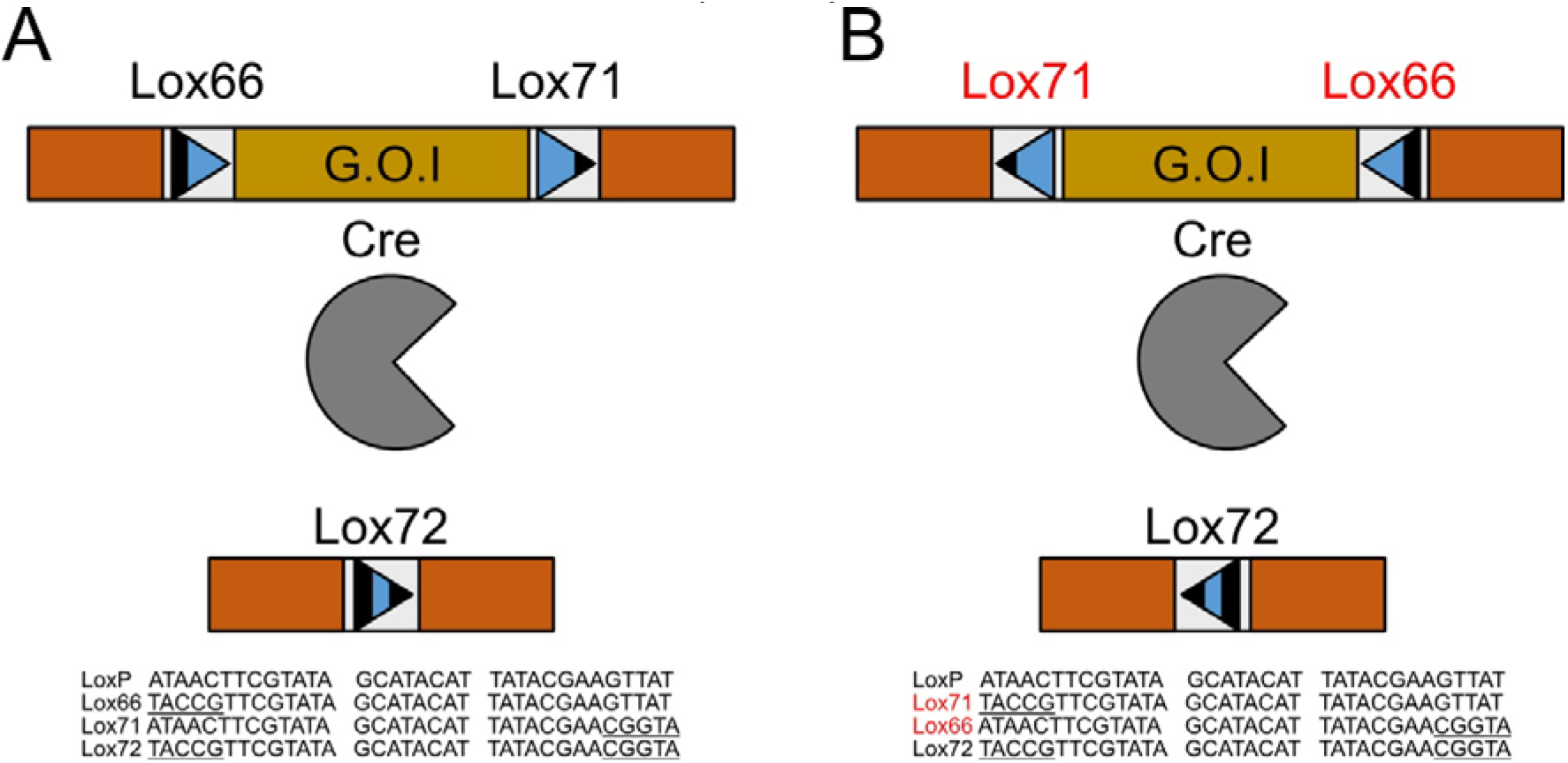
Simplified experimental schematic, modified from Figure 1 of Weng et al (42). (A) Simplification of the original Figure 1 from Weng et al., (42), showing the directionality and labels attributed to the lox sites used, and their specific sequences. (B) Our modified figure, showing the true annotations and orientations of the construct, with changes in red.

The use of incorrect lox sites has the potential to severely undermine experimental design. If the identity of a lox site is incorrectly determined, then a deletion that aimed to create an inactive lox72 could instead create an active loxP. We have shown that the action of the cre upon a single active lox site within the genome can have severe consequences for the cell, showing a lethal phenotype in *M. pneumoniae*.

The cause of this lethality is currently unknown, however we currently have two main hypotheses. The first, is that the Cre attempts the reaction despite the second lox site not being present. The cleaved leading strand then has no receptor, and thus a nick is formed. If the lagging strand is also cut, then this will form a double stranded break in the DNA, which is highly lethal in most bacteria (62).

The second hypothesis is that the Cre becomes irreversibly bound to the DNA, as the reaction stalls due to lack of components. This may leave the Cre bound to the DNA during the strand cleavage reactions, and thus not only create nicks and potential double strand breaks, but also become a hindrance to DNA transcription, translation and replication machinery.

Regardless of the cause, it is apparent that this action has applications in engineering purposes, as either a kill switch mechanism or selection against cells that have undergone unwanted DNA inversions. This was demonstrated by Shaw et al., (in preparation) as a highly effective counter-selection system to remove unwanted inversions when lox sites were added randomly to the genome of *M. pneumoniae*, leaving only cells that had undergone a deletion and produced an inactive lox72 site, as cells that contained inversions which produced active lox sites were killed.

## Conclusions

Of the 21 articles that unambiguously showed the correct lox site annotations, 18/21 used the same lox orientation as the original paper by Albert et al. (8), with the lox directionality reading from right to left (8,12,17–23,25–34). In contrast, 5 of the 8 papers with mis-labelled lox sites portray them from left to right (35,37,39–41). It seems apparent that there is potential for the mis-annotation of these sequences, and that this could stem from incorrectly assuming either lox site directionality, or by accidentally attributing the wrong name to the correct site.

While the cases identified here are benign, with miss-annotations not affecting the outcomes of the experiments, the prevalence of these errors is worrying. Attempts at replication, or researches using these maps or constructs as the basis for their own experiments may inadvertently create incompatible designs of vectors, and thus cause delays and errors in future experiments. We have demonstrated that the action of the Cre on a single lox site can have a lethal effect, and therefore the authors believe that special care should be taken to properly attribute the correct names to lox sites.

As such, it seems prudent to recommend that researchers continue using the annotations provided by Albert et al (8), repeated below, as this is the form taken by the majority of the correct cases, and unused by the majority of the incorrect ones. We also recommend the avoidance of the left or right element notation, as it is too ambiguous and dependent on orientation. If the lox site is written from right to left, then the LE mutant would be the lox71, however if the lox site is written left to right, then the LE becomes lox66. Therefore in the aim of standardisation, we recommend following the original notation and designing lox sites from right to left whenever possible:

Lox66: 5’ - ATAACTTCGTATA GCATACAT TATACGAAcggta - 3’
Lox71: 5’ - taccgTTCGTATA GCATACAT TATACGAAGTTAT - 3’

By adopting a more standardised nomenclature, hopefully the internal reproducibility and accuracy of the reporting of lox sites, and errors or confusion caused by the usage of incorrect sites can be minimised.

## Supplementary Tables

**Table S1:**
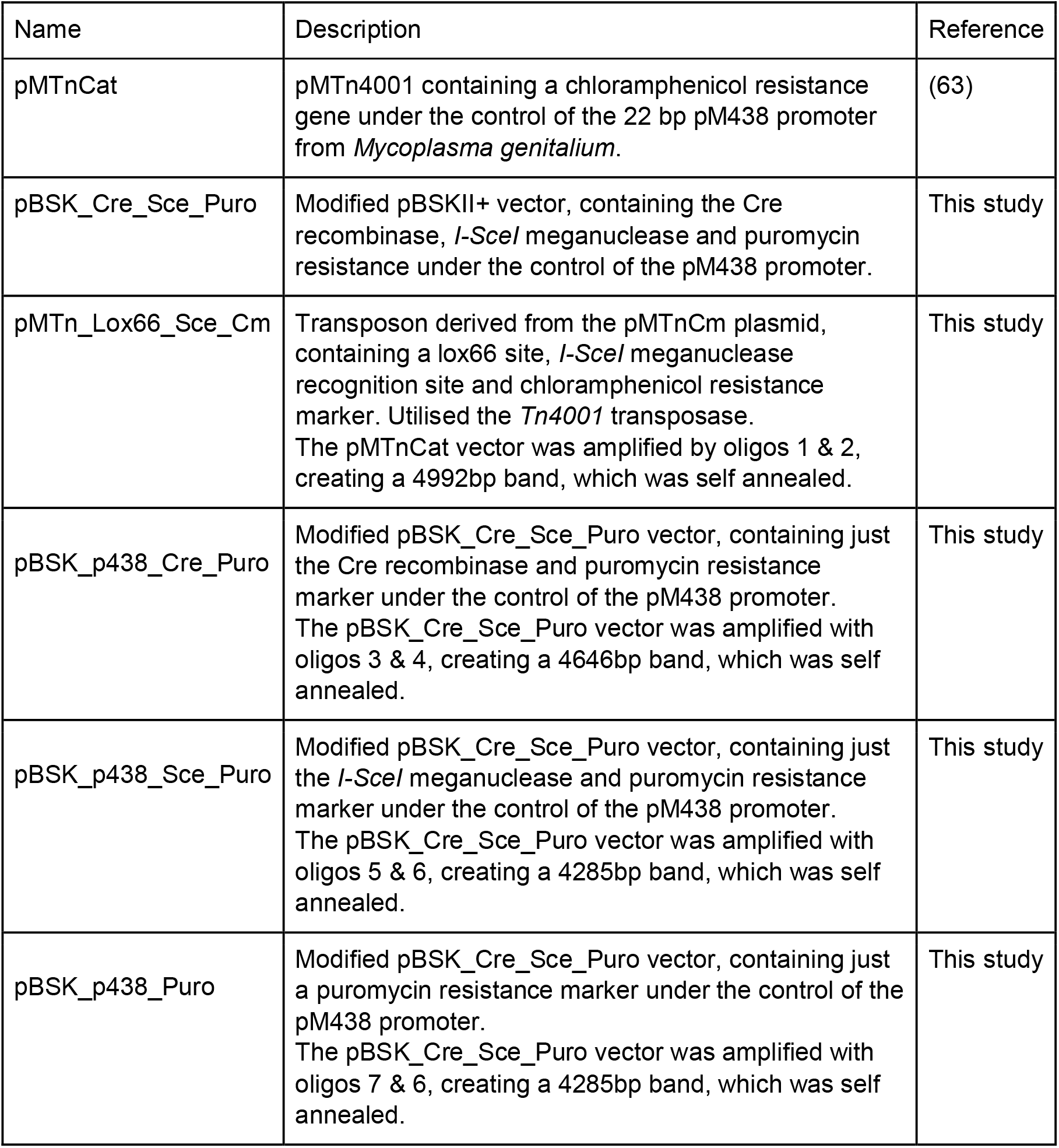
List of all plasmids used in this study.

**Table S2:**
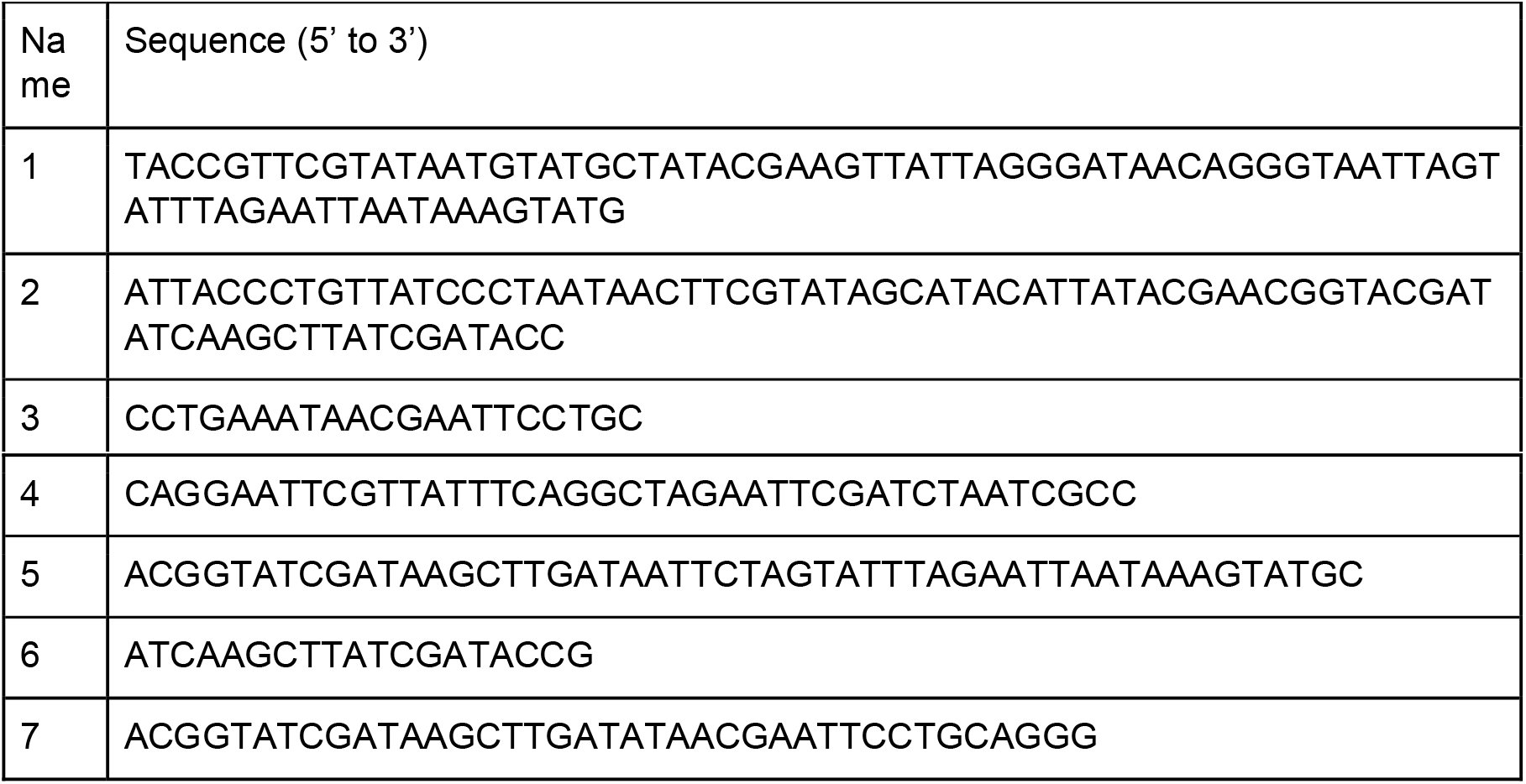
List of oligos used in this study

## Author Contributions

Daniel Shaw:

Conceptualization
Data curation
Formal analysis
Investigation
Methodology
Validation
Visualization
Writing – original draft
Writing – review & editing

Luis Serrano:

Conceptualization
Funding acquisition
Resources
Supervision
Writing – review & editing

Maria Lluch-Senar:

Conceptualization
Supervision
Writing – review & editing

